# Long non-coding RNAs contribute to DNA damage resistance in *Arabidopsis thaliana*

**DOI:** 10.1101/2023.03.20.533408

**Authors:** Nathalie Durut, Aleksandra E. Kornienko, Heiko A. Schmidt, Nicole Lettner, Mattia Donà, Magnus Nordborg, Ortrun Mittelsten Scheid

**Affiliations:** Gregor Mendel Institute, Austrian Academy of Sciences, Vienna BioCenter (VBC), Vienna, Austria; Center for Integrative Bioinformatics Vienna (CIBIV), Max Perutz Labs, University of Vienna and Medical University of Vienna, Vienna BioCenter (VBC), Vienna, Austria

**Keywords:** Long non-coding RNA, DNA damage, DNA repair, double strand break, plant, Arabidopsis, Brassicaceae

## Abstract

Efficient repair of DNA lesions is essential for faithful transmission of genetic information between somatic cells and for genome integrity across generations. Plants have multiple, partially redundant and overlapping DNA repair pathways, probably due to the less constricted germline and the inevitable exposure to light including higher energy wavelengths. Many proteins involved in DNA repair and their mode of actions are well described. In contrast, a role for DNA damage-associated RNA components, evident from many other organisms, is less well understood. Here, we have challenged young *Arabidopsis thaliana* plants with two different types of genotoxic stress and performed *de novo* assembly and transcriptome analysis. We identified three long non-coding RNAs (lncRNAs) that are lowly or not expressed under regular conditions but up-regulated or induced by DNA damage. To understand their potential role in DNA repair, we generated CRISPR/Cas deletion mutants and found that the absence of the lncRNAs impairs the recovery capacity of the plants from genotoxic stress. The genetic loci are highly conserved among world-wide distributed Arabidopsis accessions and within related species in the *Brassicaceae* group. Together, these results suggest that the lncRNAs have a conserved function in connection with DNA damage and provide a basis for a mechanistic analysis of their role.

## Introduction

Insight into diversity and functions of non-coding RNAs (ncRNAs) without a potential to code for more than short peptides is growing constantly. Some can be classified according to conservation of sequences and functions like tRNAs or rRNAs; others differ by sequence but form functional categories, e.g., miRNAs. In recent years, the enormous amounts of RNA sequencing data provided evidence for the existence of numerous additional RNA varieties, and for most of them, a functional taxonomy is still missing. Size is a convenient distinction, and there is a general agreement to call those above a length of 200 nt long non-coding RNAs (lncRNAs), although this coarse classification seems like a surrender facing the enormous diversity of their form and function (Mattick et al., 2023). The category comprises lncRNAs ranging from those expressed constitutively in all cell types and with well-defined roles to others present only in special cells, under exceptional conditions and so far without insight into their biological context (Mattick et al., 2023). The latter are by far the majority, and understanding their contribution to differentiation, development, growth, adaptation, or disease will be challenging and rewarding. Although the mode of lncRNA action is even less understood than their role, they can exert regulatory roles by interaction with proteins, DNA, or other RNAs, leading directly or indirectly to altered expression of protein-coding genes. It is likely that diversity of lncRNA, in numbers and function, solves the “g-value paradox” referring to the discrepancy between similar numbers of protein-coding genes and widely varying organismal complexity (Hahn and Wray, 2002; Mattick et al., 2023).

Although lncRNAs are found in all organisms, plant research has contributed substantially to confirm their biological relevance, as evident from a wealth in recent review literature (Ben Amor et al., 2009; Chen et al., 2020; Bhogireddy et al., 2021; Chekanova, 2021; Jampala et al., 2021; Wierzbicki et al., 2021; Chao et al., 2022; Ma et al., 2022; Roulé et al., 2022; Sharma et al., 2022; Zhao et al., 2022). Many reports indicate a connection of lncRNA expression with external challenges, like pathogen attack, nutrient limitation, or other abiotic stress types. The sessile lifestyle of plants might have been an evolutionary force to drive diversification of lncRNAs as regulatory elements especially in this context.

One of the stress factors for which a connection with and a role of lncRNAs was postulated or documented is DNA damage and its repair (reviewed in Fijen and Rothenberg, 2021; Guiducci and Stojic, 2021; Shaw and Gullerova, 2021; Zhu et al., 2022; Yu et al., 2023). Most of these reports are about mammalian cells, very prominently in connection with genetic instability in cancer cells. Beside some diversification in DNA repair, basic principles are shared between plants, fungi, and animals. It is therefore likely that DNA damage repair in plants could also include RNA components. The dependence of plants on light is intrinsically connected with their exposure to the UV part of the spectrum, causing several types of DNA damage than can result in deleterious mutations. Besides other protective means, e.g., producing absorbing pigments or adjusting leaf orientation, plants have several pathways for efficient DNA damage repair and maintenance of genome integrity, and numerous proteins of this portfolio are well characterized (Bray and West, 2005; Balestrazzi et al., 2011; Gill et al., 2015; Manova and Gruszka, 2015; Nisa et al., 2019; Hacker et al., 2020; Casati and Gomez, 2021). Insight into a potential involvement of lncRNAs in plant DNA repair is emerging (reviewed in Durut and Mittelsten Scheid, 2019), mainly connected with the most dangerous type of DNA lesions by double-strand breaks (DSBs), but so far not well documented.

Here, we describe the screen for lncRNAs in the model plant *Arabidopsis thaliana* that are induced upon the generation of DNA double strand breaks by genotoxic stress. Among several candidates, we characterized three of them in detail and provide evidence that their loss affects the ability of plants to recover from DNA damage. This important functional role is further supported by their sequence conservation between accessions of multiple origins and within the Brassicaceae.

## Results

### Genome-wide identification of lncRNAs in response to DNA damage

To study whether DNA damage would relate to lncRNAs in plants, we exposed 15-day-old Arabidopsis seedlings to genotoxic conditions that would create several randomly distributed lesions in genomic DNA. We applied two mechanistically different treatments: either zeocin, a drug that chemically generates both single and double strand breaks (DSBs), or UV-C irradiation, which induced the formation of pyrimidine dimers and other photoproducts, as well as reactive oxygen species (ROS) which can result in DSBs. Treated and non-treated samples (mock) were used to prepare RNA. This was depleted from ribosomal RNAs and used to generate strand-specific libraries which were Illumina-sequenced in the 50 bp paired-end mode (Figure 1 A).

**Figure 1:**
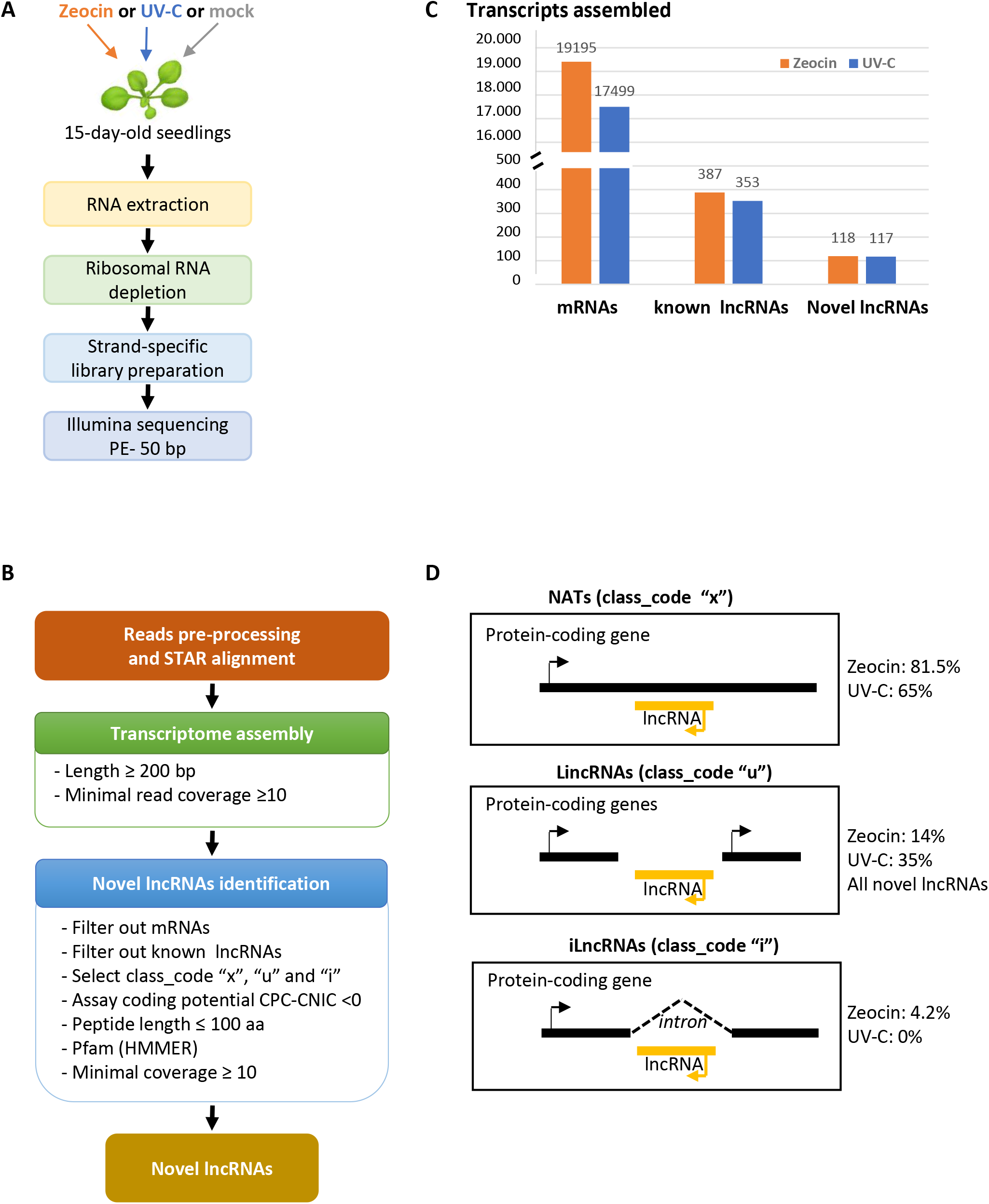
Identification of lncRNAs upon DNA damage induction. (A) Processing of zeocin-or UV-C-treated plant material for RNA sequencing. Libraries were prepared from 5 (zeocin) or 3 (UV-C) biological replicates. (B) Processing of sequence data for transcriptome assembly and lncRNA identification. (C) Transcript assembly in libraries of zeocin-or UV-C-treated samples. mRNAs and known lncRNAs were counted if present in the annotation of the reference genome Araport 11. Novel lncRNAs refer to newly identified transcribed regions. (D) Classification of lncRNAs according to their position in or between protein-coding genes.

From 5 (zeocin) and 3 (UV) independent experiments, all including mock-treated controls, we obtained a total of 505 and 255 million reads, respectively (Table S 1). Trimmed reads were aligned to the Arabidopsis reference genome (TAIR10) and assembled into transcriptomes including both mock and treated conditions. This resulted in 20,460 (mock/zeocin) and 18,535 (mock/UV) unique transcripts, respectively. Among these transcripts, 19,195 and 17,499 were mRNAs of protein-coding genes (Araport 11), and 387 and 353 were lncRNAs (annotated as either lncRNAs, natural antisense transcripts (NATs), novel transcribed regions, or other RNAs). After multiple filtering steps (Figure 1 B), we identified 118 and 117 novel putative lncRNAs (Figure 1 C). According to their genomic positions, they were classified as NATs (81.5% and 65%), intergenic lncRNAs (lincRNAs) (14% and 35%), and intronic lncRNAs (ilncRNAs) (4.2% and 0%) in zeocin- and UV-treated samples, respectively (Figure 1 D).

We characterized the features of the novel lncRNAs, including their average size, the number of exons, and their expression level and compared them with those for protein-coding transcripts (mRNAs). With a mean length of 887 nt, lncRNAs in zeocin-treated samples were on average shorter than mRNAs with a mean of 1660 nt; 710 nt versus 1685 nt for UV-treated samples (Figure S1 A). In addition, lncRNAs had significantly less exons (mean 1.3 and 1.5 exons, respectively) than mRNAs (mean ~ 5 exons) and lower expression levels (Figure S 1 B and S 1 C), which is in agreement with previous studies (Zhao et al., 2018).

### Identification of lncRNA genes responding to DNA damage

To identify RNAs with a specific response to DNA damage, we compared the transcriptome from mock-treated plants with those subjected to DNA damage. In total, we identified 29 and 194 differentially expressed (LFC >1.5) lncRNAs (both annotated and novel lncRNAs) in zeocin- and UV-treated samples, respectively, in addition to 473 and 2603 differentially expressed protein-coding genes (Figure 2 A, B). The analysis validated the induction of DNA damage, by the apparent up-regulation of DNA repair marker genes like *BRCA1, RAD51*, and *PARP2* upon zeocin treatment (Doutriaux et al., 1998; Doucet-Chabeaud et al., 2001; Lafarge and Montané, 2003) and *GST1, MC8*, and *CAT2* (Rentel and Knight, 2004; Vanderauwera et al., 2011; Tang et al., 2016) in response to UV stress (Figure S 2 A), as well as a GO term enrichment for DNA repair and recombination (zeocin) or general stress response (UV) (Figure S 2 B). By comparing the two datasets, we found in total 149 genes that are differentially expressed compared to the mock controls and are shared by both treatments, including lncRNAs (Figure 2 B). Two of those differentially expressed lncRNAs are significantly up-regulated after both treatments. A third lncRNA just below the significance threshold in the UV RNA-seq data was included for further analysis. The up-regulation evident from the RNA-seq data was further validated by quantitative RT-PCR analysis in zeocin and/or UV-C treated samples (Figure 2 C). All three lncRNAs loci are already annotated in the reference genome (Araport 11), and we named them lncRNA B (AT4G07235), lncRNA C (AT4G09215), and lncRNA D (AT3G00800). Their genes are located on the arms of chromosomes 3 and 4 (Figure 3 A). We determined the 5’ and 3’ ends of the transcripts by RACE-PCR, resulting in lengths of 391 nt, 443 nt, and 361 nt for lncRNA B, C, and D, respectively, with minor deviations from the annotation (Figure 3 B). The genomic loci of all three lncRNAs were enriched in zeocin-treated material after immunoprecipitation of RNA polymerase II, indicating that they are products of the same transcription process generating mRNAs (Figure 3 C). Successful amplification with oligo(dT) primers (Figure 2 C and 3 D) confirms that they are polyadenylated. None of the three lncRNAs has a protein-coding potential for more than 100 amino acids. lncRNAs C and D could be translated into short peptide sequences, but none of them has been found in a data set from a proteomic analysis of plant material after DNA damage treatment (Roitinger et al., 2015). The specific association of lncRNA B, C, and D with DNA damaging conditions is further supported by the observation that their induction by zeocin treatment is significantly reduced in the background of the *atm* mutant, lacking one of the kinases signaling DNA damage to repair pathways (Garcia et al., 2003) (Figure 3 D). ATM dependency for induction was also confirmed for five additional assembled, but previously not annotated or identified lncRNAs that are also differentially expressed upon zeocin treatment but not further studied here (Supplemental Figure 3). Taken together, the induction of otherwise not or lowly expressed lncRNAs by genotoxic treatments creating random lesions, and its dependence on DNA damage perception, suggest a specific response and a functional role for them in dealing with DNA repair.

**Figure 2:**
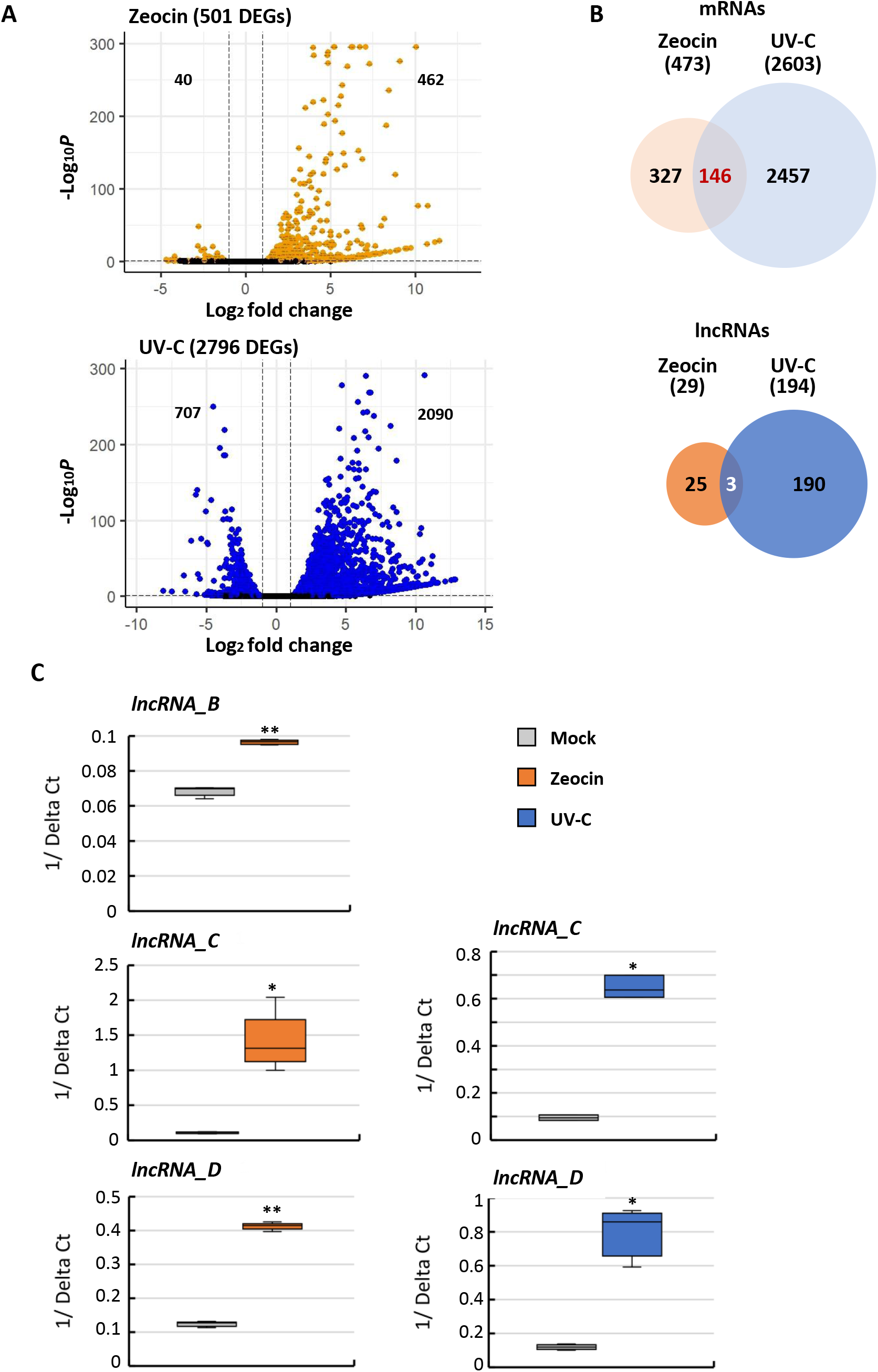
Comparison of differentially expressed genes. (A) Differentially expressed genes after treatment with zeocin (upper panel, 20880 variables) or UV-C (lower panel, 22911 variables). Numbers of down-or up-regulated genes above threshold (orange or blue versus green dots) are indicated. (B) Venn diagrams for overlap between mRNAs (upper) or lncRNAs (lower) differentially expressed between treated and mock-treated zeocin-or UV-C samples. (C) RT-qPCR validation of differential expression of lncRNAs induced by zeocin- and UV-C-treatment Error bars indicate standard deviation of 3 biological replicates (*Welcher test* **p-value <0.01, *p-value <0.05).

**Figure 3:**
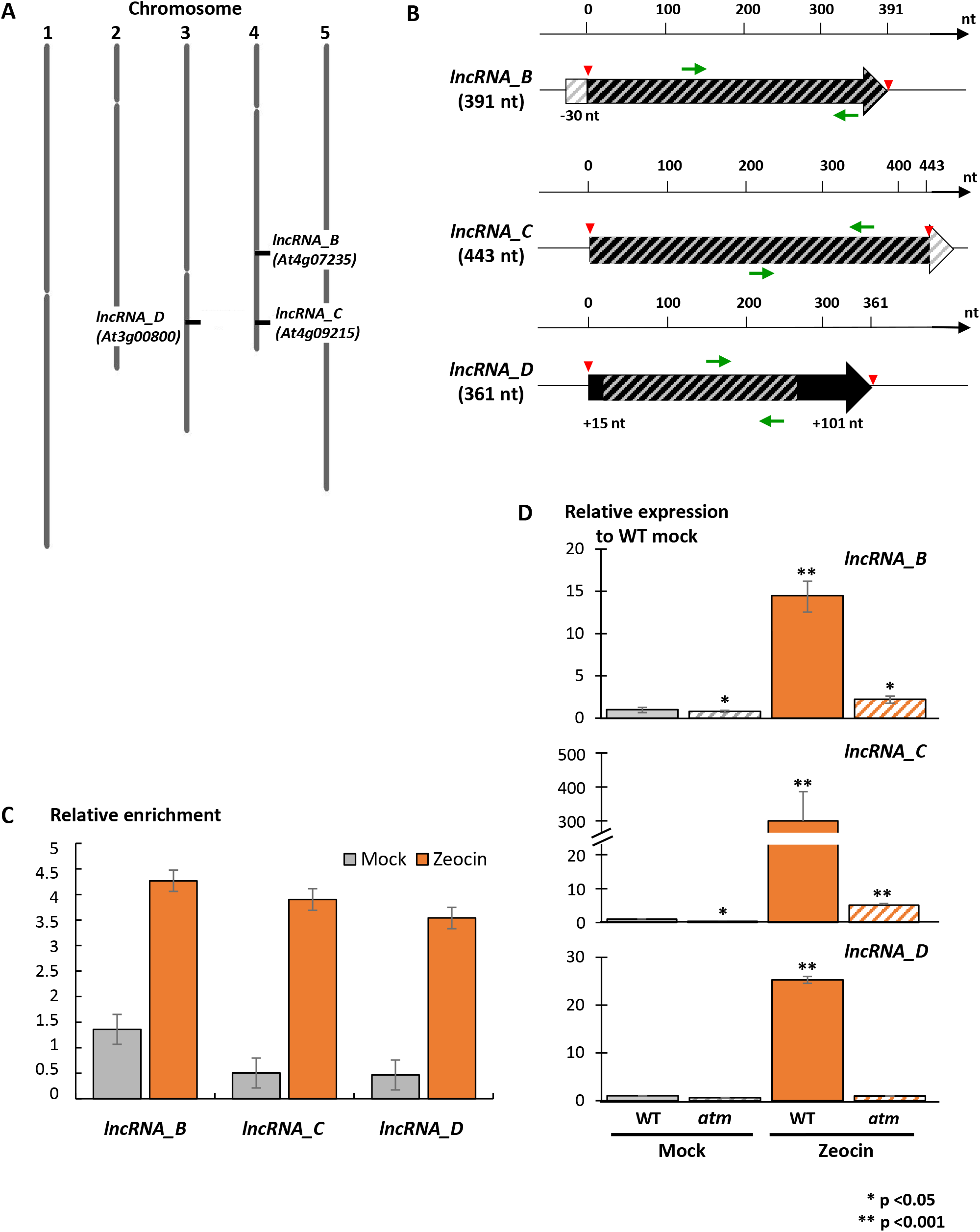
Characterization of DNA-damage-induced lncRNAs. (A) Location of the genes encoding lncRNA B, C, and D on chromosome 3 and 4 of *Arabidopsis thaliana*. (B) Scheme of the genes encoding lncRNA B, C, and D. Black arrows represent lncRNA transcripts confirmed by RACE-PCR; grey stripes represent the annotation in Araport11. Green arrows indicate the position of the two primers used for 5’ or 3’RACE-PCR. Red triangles indicate the 5’ and the 3’ends identified by RACE-PCR. (C) RT-qPCR with specific primers for lncRNAs B, C, or D on chromatin samples immunoprecipitated with a PolII antibody recognizing the CTD domain, from mock-treated or zeocin-treated samples. (D) Expression of lncRNAs B, C, or D in WT or *atm* mutant in mock-or zeocin-treated samples, normalized to a constitutively expressed actin gene. Data were normalized to the values in WT mock samples. Error bars indicate standard deviation of 3 biological replicates (*Welcher test* **p-value <0.01, *p-value <0.05).

### Determining DNA damage sensitivity in mutants lacking lncRNA genes

To assay the role of lncRNAs B, C, and D in the context of DNA damage, we decided to challenge loss-of-function mutants with genotoxic stress. As there were no suitable mutants for any of the three genes available in the stock center collections, we generated deletion mutants with the CRISPR gene editing approach. We designed sgRNAs aiming for a complete deletion of the corresponding genes by designing sgRNAs outside of the annotated region and succeeded in generating homozygous deletions for all three loci. Plants with these genotypes were slightly delayed in growth but had an otherwise regular morphology (Supplemental Figure 4). By northern blots with probes covering the full length of the genes, we confirmed that no sequences homologous to the lncRNA transcripts were detectable in the mutant plants, neither in mock nor in zeocin-treated plants (Figure 4 A).

**Figure 4:**
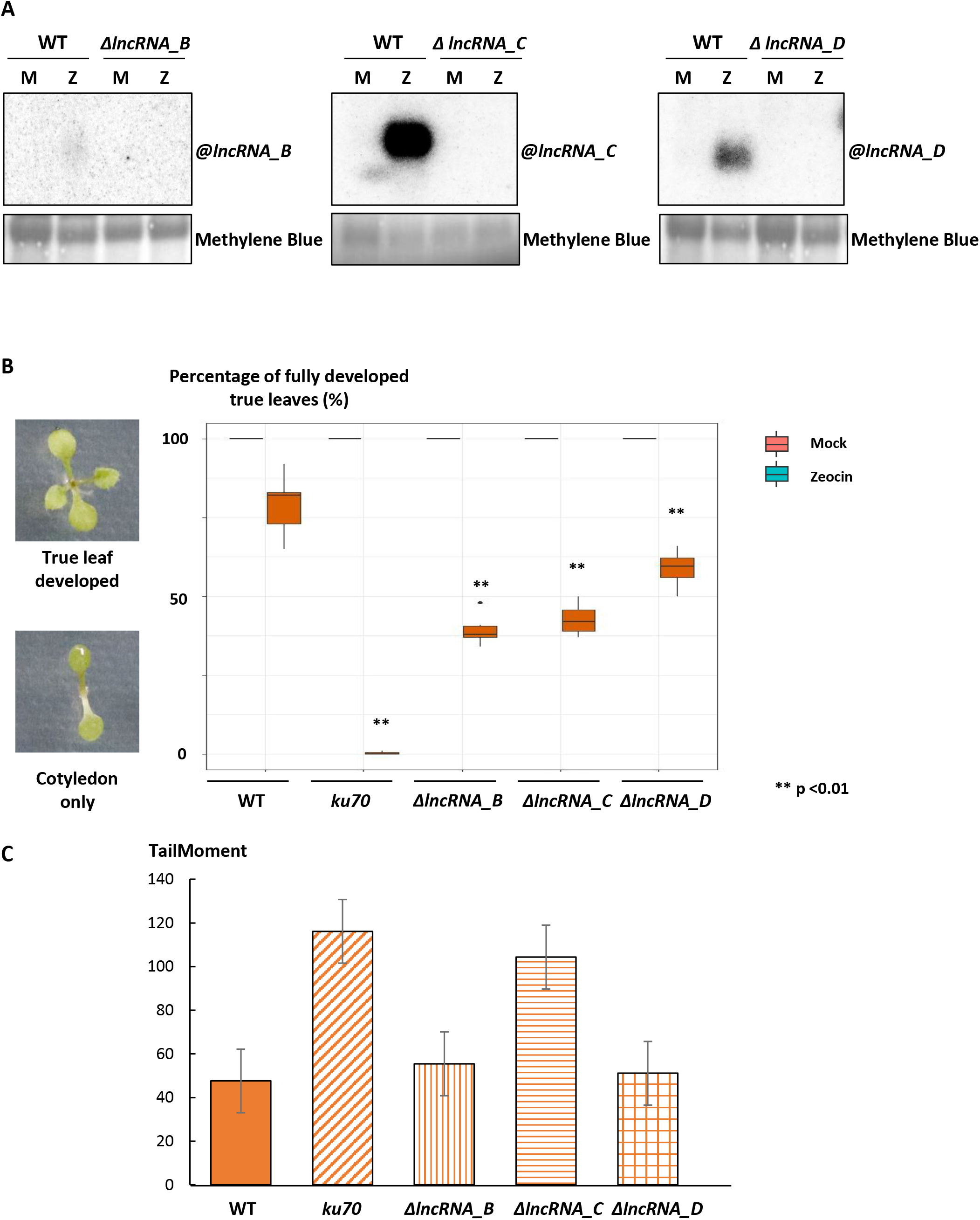
Characterization of deletion mutants. (A) Norther blots with total RNA probed with the radioactively labelled amplicons for lncRNAs B, C, or D, in WT or plants in which the lncRNA gene had been deleted by CRISPR-Cas9 mutations. M: mock-treated; Z: zeocin-treated; Methylene Blue: loading control. (B) True-leaf assay for plant sensitivity against DNA damage. Left: seedlings resistant to zeocin can grow and develop true leaves; sensitive seedlings are arrested after cotyledons have unfolded. Right: Resistance ratio in WT, *ku70* as a known sensitive repair mutant, or the deletion mutants lacking lncRNA B, C, or D (p-values according to Mann-Whitney-test). (C) Comet assay for plant sensitivity against DNA damage.

To test whether the deletion mutants would be more sensitive to DNA damage than the wild type, we applied the well-established true-leaf assay (Rosa and Mittelsten Scheid, 2014). In brief, seeds are surface-sterilized and sown on solid growth medium containing a defined dose of zeocin, so that the developing seedlings are exposed to a limited dose of genotoxic stress. Later, they are scored for development of true leaves, indicating the potential to repair DNA damage and continue growth (Figure 4 B). Quantification of the ratio between seedlings with true leaves and all exposed seedlings reveals good recovery of the wild type, in contrast to strongly impaired recovery of *ku70*, a mutant with a defect in DNA repair by non-homologous end joining (Riha et al., 2002). Recovery of all three lncRNA deletion mutants was also reduced, not as drastically as the *ku70* mutant but significantly different from the wild type (Figure 4 B). We also applied the comet assay, an independent quantitative test for DNA damage repair capacity. Here, nuclei of mock-or zeocin-treated plant material are embedded into agarose and subjected to electrophoresis. The amount of DNA fragments pulled into the direction of the anode, forming a comet tail, indicates the degree of non-repaired DNA (Menke et al., 2001). In this assay, the mutant lacking lncRNA C shows a clear repair deficiency, similar to that in *ku70*, whereas the difference to the wild type is not significant for the lncRNAs B and D mutant (Figure 4 C).

### Conservation of lncRNA genes within Arabidopsis accessions

To explore if lncRNAs B, C, and D would be induced by genotoxic stress beyond the reference accession Col-0, we exposed seedlings of five other accessions to zeocin and determined expression of the lncRNAs by quantitative RT-PCR. While there was measurable induction compared to mock controls for all three lncRNAs in most accessions, there were striking and reproducible differences in the degree of induction (Figure 5 A). This stimulated us to explore the sequence diversity at the genomic loci within multiple Arabidopsis accessions originating from different habitats around the Northern hemisphere (Kawakatsu et al., 2016).

**Figure 5:**
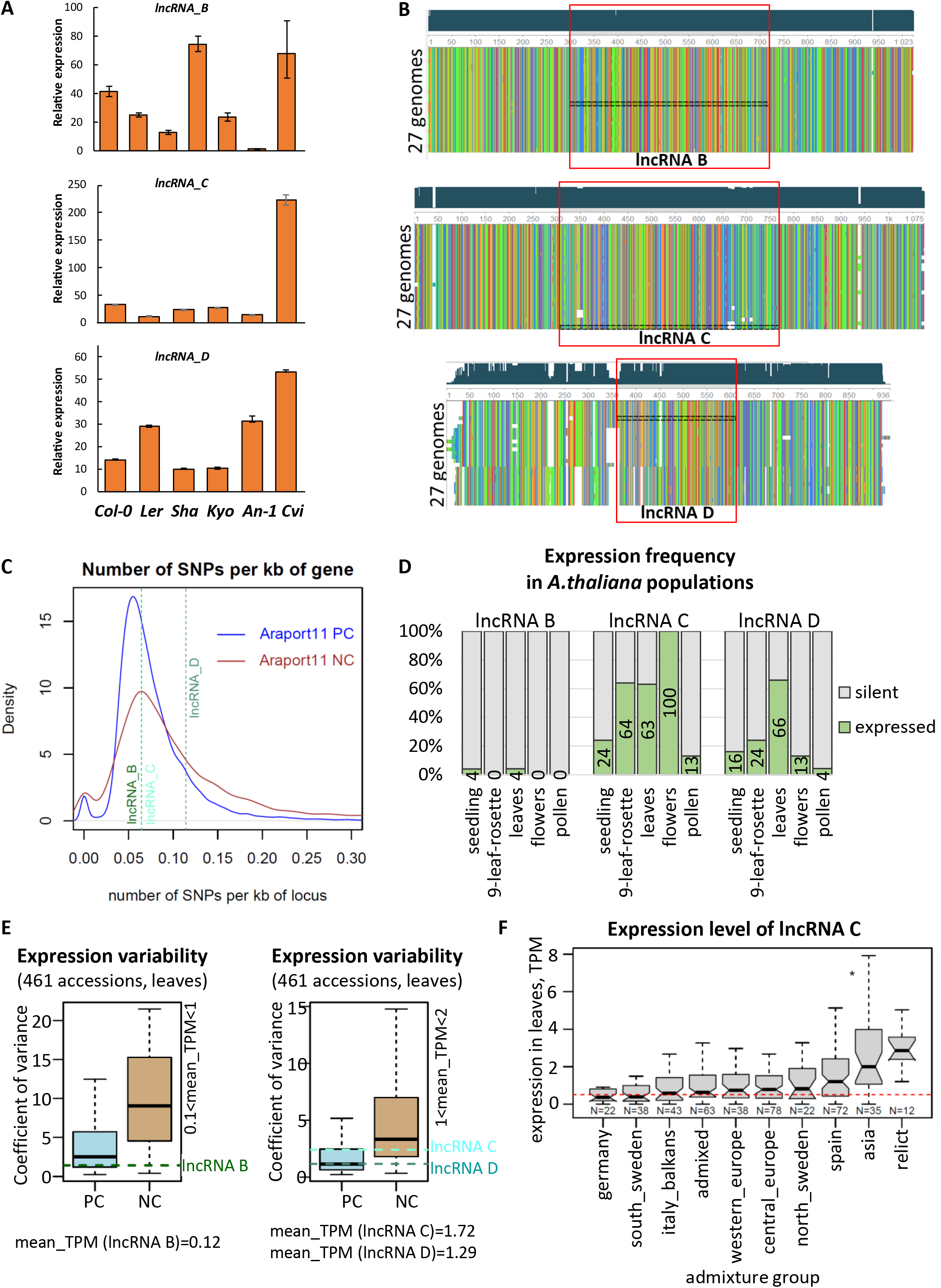
Conservation of genes for damage-associated lncRNAs B, C, or D. (A) Relative expression level of lncRNAs B, C and D in Col-0 and five non-reference accessions upon exposure to zeocin. The relative expression is the ratio between treated samples and mock controls for each accession. Error bars represent standard deviation across 3 replicates. (B) Multiple alignments of lncRNAs B, C and D loci and their flanking 300 bp regions identified in the full genomes of 27 *A. thaliana* accessions. The red boxes mark the regions corresponding to the lncRNA transcript. The narrow black box indicates the Col-0 reference accession. The multiple alignments are sorted (descending) by length. (C) Distribution of the number of SNPs per 1 kb for Araport11-annotated protein-coding genes (blue line) and non-coding genes (red line). Dashed vertical lines show the exact number of SNPs per 1 kb for lncRNA B (dark green), lncRNA C (light aquamarine), and lncRNA D (dark aquamarine). The number of SNPs is calculated according to the SNP calling from 1135 natural *A. thaliana* accessions (https://1001genomes.org/accessions.html). (D) Percent of *A. thaliana* natural accessions that express (TPM>0.5) lncRNAs B, C or D. The ratios were calculated from RNA-seq data from seedlings, rosettes at the 9-leaf stage from 25 accessions, flowers and pollen from 23 accessions (Kornienko et al., 2023), and leaves from mature pre-bolting rosettes from 461 accessions (Kawakatsu et al., 2016). (E) Expression variability across 461 accessions ((Kawakatsu et al., 2016) for lowly expressed lncRNA B (left) and moderately expressed lncRNA C and D (right), compared to that for lowly expressed Araport11 protein-coding (PC) and non-protein-coding (NC) genes. The precise level of the expression variability of lncRNAs B, C and D is indicated with horizontal dashed lines. Data source as in Figure 5 D. (F) Expression levels of lncRNA C in accessions of different geographic origin defined by admixture groups. The red dashed horizontal line indicates expression cut-off (TPM=0.5). Data source as in Figure 5 D. The admixture group of each accession was determined based on genetic similarity (http://1001genomes.github.io/admixture-map/).

The analysis of the SNP data from 1135 accessions (https://doi.org/10.1016/j.cell.2016.05.063) showed that lncRNAs B and C had a similar number of SNPs per kb as many other lncRNAs annotated in Araport11, while lncRNA D showed much less conservation (Figure 5 B). We then analyzed 26 full genome assemblies of non-reference *A. thaliana* accessions (provided by the Nordborg lab, GMI, Austria) for evidence of copy number differences and structural variation in the three lncRNA loci. In agreement with the SNP analysis, lncRNA D showed the highest variability, with short sequences missing in some accessions, particularly in the upstream region. lncRNAs B and C are highly conserved (Figure 5 C). All three lncRNA genes are present in only one copy in every of the 27 accessions, and they do not contain sequences related to transposable elements.

Analyzing expression data from multiple accessions (Kawakatsu et al., 2016; Kornienko et al., 2023) generated from soil-grown plants without genotoxic stress indicated absence of transcripts of all three lncRNAs in the reference accession Col-0 seedlings (Supplemental Figure 5 A) but expression in seedlings from some other accessions under the same conditions. This is rare for lncRNA B but most common for lncRNA C (Figure 5 D). There are tissue-specific differences, as lncRNA C is detectable in flowers of all accessions (including Col-0, Supplemental Figure 5 A), while lncRNA D is more often expressed in mature leaves (Figure 5 D, Supplemental Figure 5 B). Compared to most annotated lncRNAs, the expression variability of lncRNAs B, C and D in leaves across 461 accessions is lower (Figure 5 E), with lncRNA C being slightly more variable than lncRNAs B and D. This expression variability for lncRNA C is more pronounced when considering geographic patterns: high in Asian accessions and relict accessions originating from ancestral habitats, but low in German accessions that include Col-0 (http://1001genomes.github.io/admixture-map/ (Figure 5 F).

### Conservation and phylogenetic analysis of lncRNA genes among other Brassicaceae

As the three lncRNAs are conserved within the different *Arabidopsis* accessions, we asked if these lncRNAs have conserved orthologs in other species beyond *Arabidopsis thaliana*. Furthermore, we were interested in their taxonomic distribution. Collecting sequences homologous to the lncRNAs B, C and D from full genomes using BLAST in different sequence databases revealed significant hits only inside the *Brassicaceae*. Accordingly, we performed phylogenetic analysis within *Brassicacean* species with available reference genomes.

The phylogenetic trees (Figure 6 A-C) show that lncRNA B, C and D are well represented in Brassicaceae Lineage I that contains *Arabidopsis thaliana*. Representation differs in Lineage II, which includes Brassica species like rapeseed and the cabbages. lncRNAs B and C are well represented there (Figure 6 A+B), whereas lncRNA D was found once (*Thlaspi arvense*) in Lineage II (Figure 6 C).

**Figure 6:**
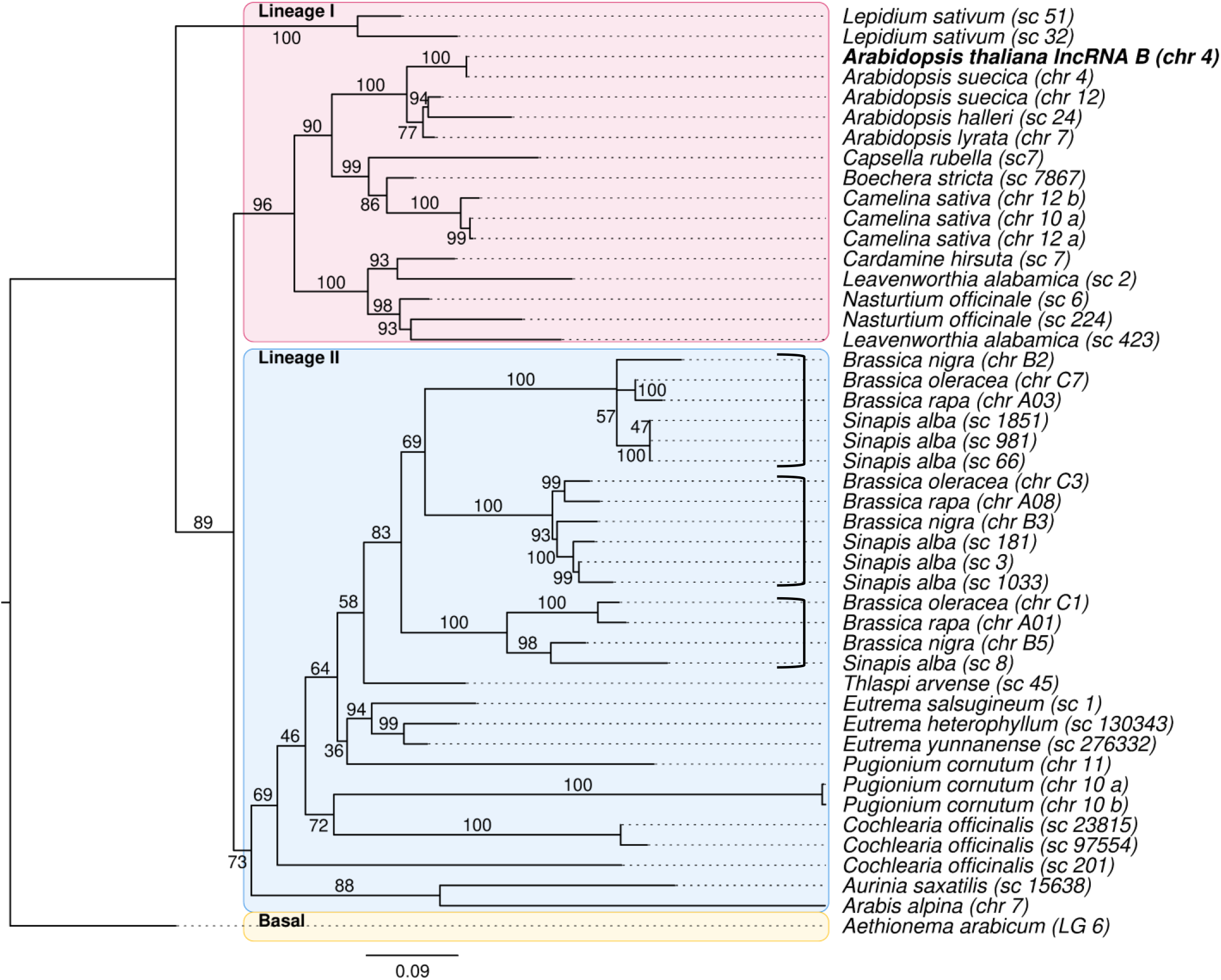
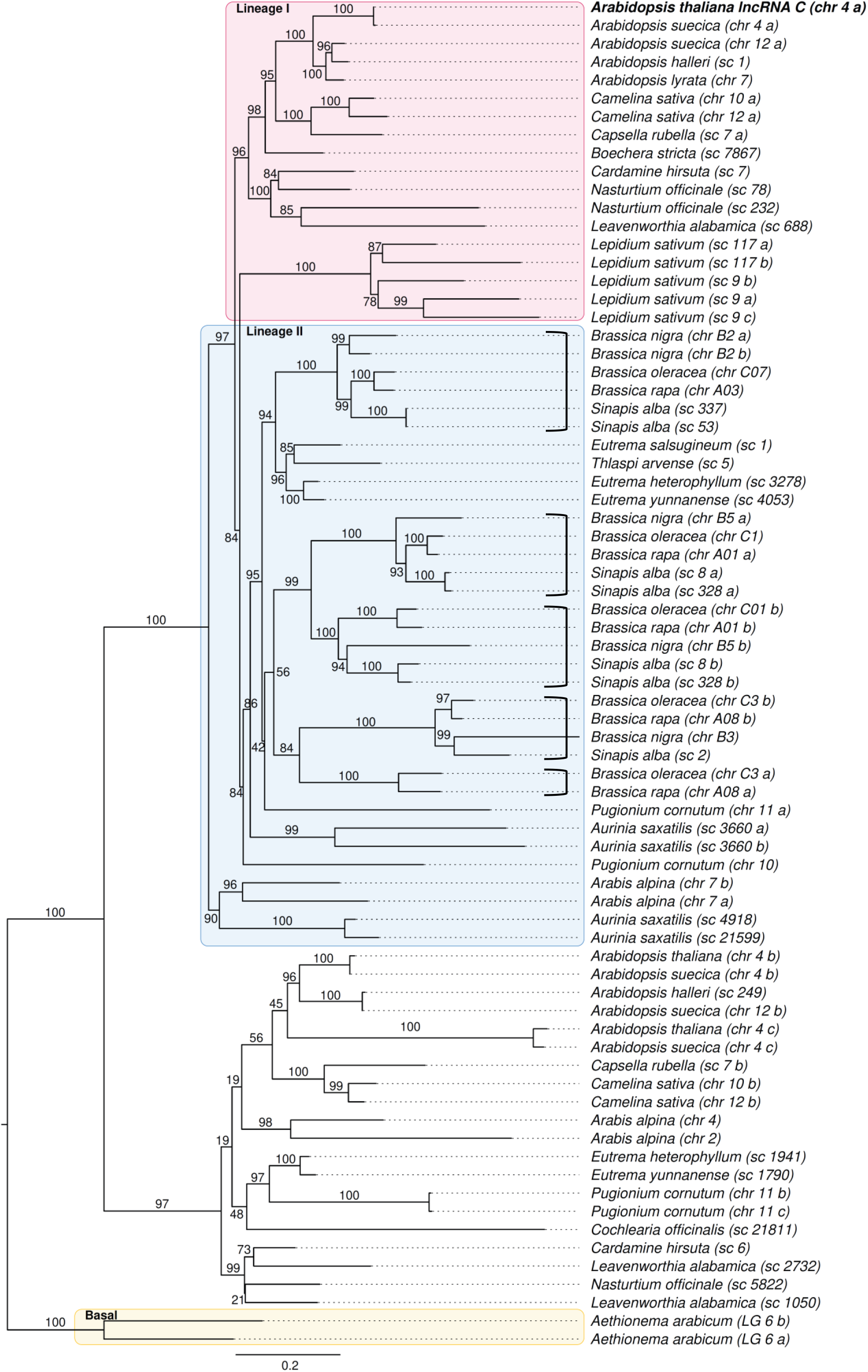
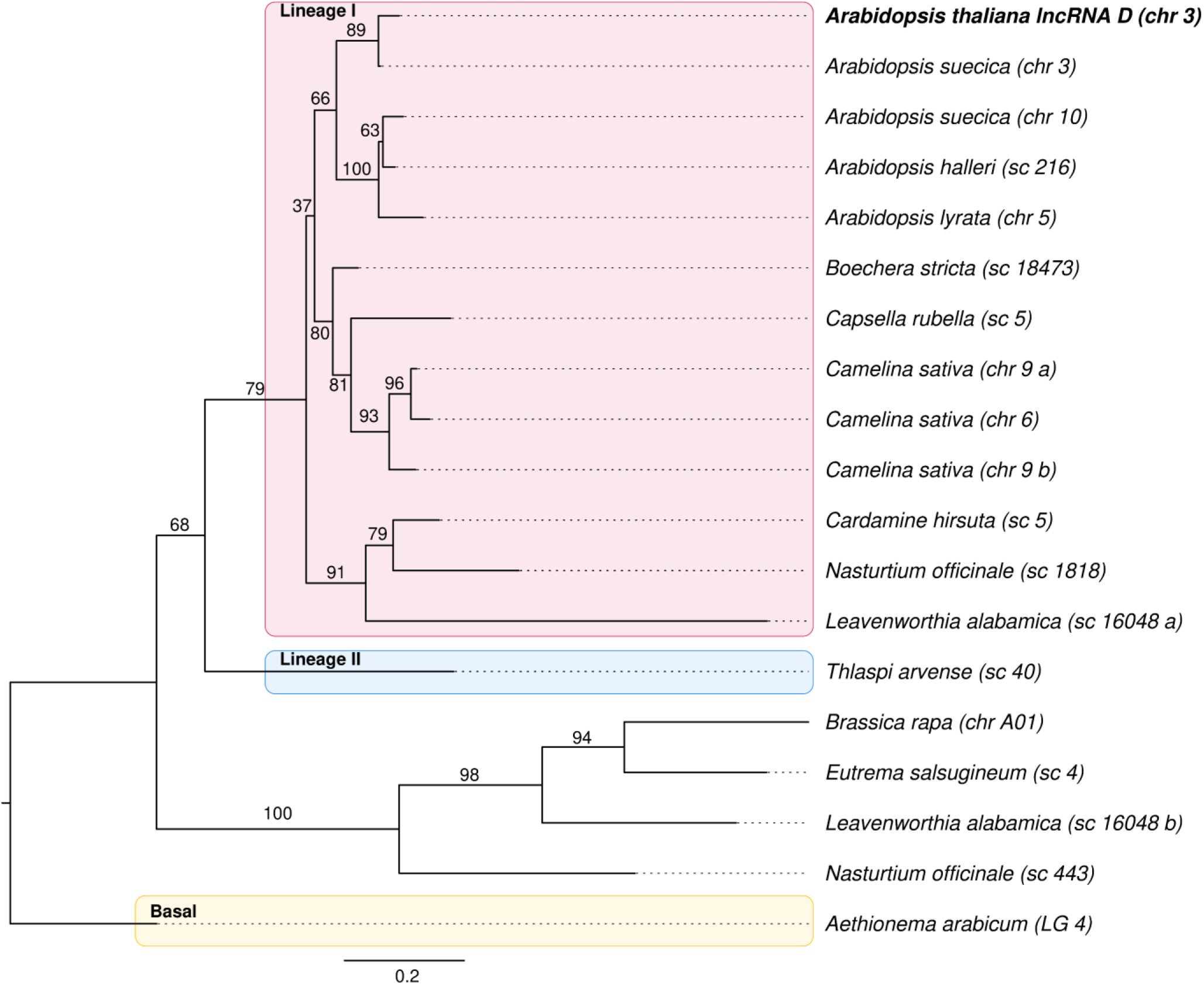
Conservation of genes for damage-associated lncRNAs B, C, or D among the Brassicaceae. Maximum Likelihood phylogenetic trees of homologous sequences to (A) lncRNA B (At4g07235), (B) lncRNA C (At4g09215), and (C) lncRNA D (At3g00800), obtained from *Brassicacean* species. The support values at the branches have been obtained from 10000 UFboot samples. The trees are rooted at the most basal species in the dataset, *Aethionema arabicum*. The sequence names denote the species as well as the chromosome (chr), supercontig (sc), or linkage group (LG) of the respective genome. If more than one sequences originate from the same chromosome, letters a, b, c etc. were appended to the sequence name. The boxes denote Brassicaceae Lineages I and II as well as the basal lineages.

Remarkably, lncRNA copy numbers show high evolutionary dynamics, which often seem species-specific. For lncRNA B, we find two copies in *Lepidium sativum* and *Nasturtium officinale*, three copies in *Camelina sativa, Pugionium cornutum* and *Cochlearia officinalis*, while lncRNA C is present in two copies in *C. sativa, N. officinale*, and *Arabis alpina*, two pairs of two in *Aurinia saxatilis* and even five copies in the genome of *L. sativum*. Also, lncRNA D is present with three copies in *C. sativa*.

A closer look into the Brassica/Sinapis group reveals clusters of copies (3 in lncRNA B and 5 in lncRNA C) that suggest duplications at the common ancestor, followed by additional duplications in Sinapis which now exhibits 7 copies of lncRNA B as well as C (Figure 6 A+B). We could not reconstruct the exact order of duplications because individual branches in the trees have only small support values. Since the duplications are either lineage-specific or happened after the split between Lineage I and II, all resulting paralogous copies are co-orthologs (according to Koonin, 2005) to the reference lncRNAs in Arabidopsis, originating from duplication of one ortholog ancestor after speciation.

## Discussion

We confirmed that, like other organisms, plants have lncRNAs that appear in connection with genotoxic stress. We identified several transcripts, including previously non-annotated lncRNAs, that are induced by DNA damage in the model plant Arabidopsis. Finding an overlap between the candidates obtained after two different types of DNA-damaging treatments increased the probability that these lncRNAs were indeed a direct response to this type of stress, and we focused the analysis on these transcripts and their genetic loci. In the absence of genotoxic stress, the respective genes are lowly expressed, and plants lacking the corresponding genes develop normally. This suggests that these lncRNAs have a specific role in connection with DNA damage. Indeed, the deletion mutants are impaired in their ability to recover from exposure to DSB-inducing zeocin. However, this increased sensitivity is not as pronounced as for a mutant deficient in DNA repair by non-homologous end joining. This difference could indicate some redundancy between the lncRNAs and would explain why the genetic loci for the lncRNAs have not been found previously in genetic screens for impaired repair capacity. In addition, the available T-DNA mutant collections did not contain suitable alterations of the genes, and EMS mutagenesis is less likely to cause functional alterations in non-coding genes. Future analysis of double and triple mutants will address potential redundancy between the damage-associated lncRNAs.

With transcript length around 400 nt, lncRNA B, C, and D fall into the same size range as many other lncRNAs identified in different context. They also share the absence of evidence for splicing (Chen and Zhu, 2022). Beyond length and induction by DNA damage, we could not find similarity in sequence, predicted secondary structure, or upstream regulatory motifs between the three transcripts.

However, the analysis of the loci in the genomes of natural Arabidopsis accessions originating from many different habitats across the world revealed very good conservation, which strongly supports their functionality. While we have no transcription data from the different accessions after zeocin-or UV exposure, it is interesting that lncRNA D, and even more C, have detectable, but variable amounts of transcripts in several accessions without the induced damage. The preferential origin of these accessions from all over Asia does not provide a clue for an adaptation to common conditions, but higher expression in flowers could indicate some induction by the more pronounced exposure of this tissue to light, including UV wavelengths. However, natural variation of lncRNA expression could also originate indirectly, as a consequence of variation in DNA damage sensing or less densely packed chromatin, e.g., in accessions like *Cvi* (Snoek et al., 2017).

The sequence conservation of the genetic loci continues into the large group of species within the Brassicaceae. This group includes many wild plants as well as important crops and is in the focus as a source for introgression breeding towards improved stress resistance (Quezada-Martinez et al., 2021), providing a wealth of genomic information. We could establish phylogenetic trees for all three lncRNAs, though with different degrees of conservation between Lineage I (B, C, and D) and Lineage II (only B and C). The trees reveal several group-or species-specific duplications, some of them subsequential. Whether the amplification occurred together with other events or is due to specific necessity, e.g., plants like Sinapis containing aggressive secondary metabolites, needs to be investigated. As in case of the diverse Arabidopsis accessions, no transcriptome data after DNA damage induction are available for the other Brassicaceae, so that their functional role there remains to be addressed. However, considering the generally low conservation for lncRNA genes (Mattick et al., 2023), their presence and persistence in many plants related to Arabidopsis could indicate functional maintenance across evolution and will provide opportunities to challenge their role in selected species. It is likely that DNA damage-associated lncRNAs are present outside the Brassicaceae, but as no obvious orthologs were found there, their identification needs experimental approaches.

The induction of lncRNAs expression upon genotoxic stress, the DNA damage sensitivity of the deletion mutants, and the conservation of the genes support their connection with the DNA repair mechanisms, but do not provide evidence for their mode of action. It is likely that they exert their role within the nucleus, but this awaits confirmation by analysis of cytoplasmic and nuclear RNA preparations. There are multiple potential roles of lncRNAs (reviewed in Durut and Mittelsten Scheid, 2019). Many lncRNAs exert their function by binding complementary RNA or DNA. As DNA lesions by zeocin or UV are randomly distributed along the genome, it is unlikely that lncRNA bind directly at the breaks, unless small stretches of complementarity are sufficient. A preliminary analysis of the transcriptome in the mutant lacking lncRNA C with and without zeocin treatment did not reveal substantial differences in the expression of other genes. Therefore, it is more likely that the lncRNAs operate via interaction with proteins, as described for many other lncRNAs (Mattick et al., 2023). This could include tethering of proteins to certain loci, modifying signaling pathways, acting as a decoy to remove specific molecules, or contributing to subcellular structures. Recently, a report on the lncRNA COOLAIR involved in the regulation of flowering time revealed that environmental conditions resulted in alternative processing and variability in secondary structures (Yang et al., 2022). In connection with DNA repair, there is growing evidence that this involves large scale reorganization of the chromatin, e.g., by changing chromatin mobility (Meschichi et al., 2022; Meschichi and Rosa, 2023) or the formation of foci with assembly of repair factors (Hirakawa et al., 2015; Hirakawa and Matsunaga, 2019; Muñoz-Díaz and Sáez-Vásquez, 2022). It remains to be investigated whether and how lncRNAs are involved in this compartmentalization, but the candidates identified in the course of the work presented and the improved techniques to visualize RNA molecules within cells (Duncan and Rosa, 2018; Huang et al., 2020) will make these approaches possible.

## Materials and methods

### Plant materials and growth conditions

*Arabidopsis thaliana* accession Columbia (Col-0) was used in this study as wild type unless otherwise mentioned. This accession was also the parental line to generate the lncRNA deletion mutants. Accessions used in Figure 5 A were provided by the Nordborg lab. The mutants with well described DNA repair deficiencies used as controls for qRT-PCR and the true leaf assays were T-DNA insertion mutants obtained from the Nottingham Arabidopsis Stock Centre: *atm* (Sail_1223_B08) and *ku70-2* (SALK_123114C).

Plants were grown either on soil or *in vitro* on GM medium (https://www.oeaw.ac.at/gmi/research/research-groups/ortrun-mittelsten-scheid/resources/) under long day (LD) conditions (16/8 h light/dark cycles) at 21°C with standard light intensity of 120 μmol.m^-2^.sec^-1^. All seeds were surface-sterilized and kept 2 days at 4°C prior to sowing.

Before treatments with genotoxic stress, seedlings were grown for 14 days on vertically arranged plates with solidified GM plates under conditions described above. For zeocin treatment, seedlings were transferred to Petri dishes containing liquid GM with or without (mock) 200 μg/ml zeocin (stock solution 100 mg/ml, Invitrogen) and incubated for 3 h with gentle shaking, followed by washing in GM. For UV-C exposure, seedlings were exposed on the plates to 8 kJ/m^2^ UV-C light in a Stratalinker 2400 (Stratagene, La Jolla, California, US) and transferred back into growth chamber for 5 h. Control plants (mock) were placed in the Stratalinker 2400 for the same time but without UV-C light exposure. After treatment, seedlings were collected, shock-frozen in liquid nitrogen and kept at −80°C for subsequent analysis. Three biological replicates were collected for each genotype and each condition.

### True leaf assay

Seeds were plated on GM medium with or without (mock) 10 μM zeocin. Plates were kept horizontally for 10 days in standard conditions, before scoring the seedlings for those with a fully developed pair of true leaves, indicating regular growth. Seedlings with single, small and/or narrow unexpanded leaves were not considered. The ratio of zeocin-treated seedlings with true leaves was calculated in relation to those in the mock-treated batches, with three biological replicates of ~ 300 seedlings each. Statistical analyses of significance for differences were performed applying a Welch two sample t-test (α = 0.05).

### Generation of transgenic lines

All vectors for plant transformation were amplified in *E.coli* strain DH5α and plasmid preparations controlled by Sanger sequencing before being transformed into electrocompetent *Agrobacterium tumefaciens* strains GV3101. *Arabidopsis thaliana* Col-0 plants were grown in the standard conditions described above for approximately 4 weeks until they reached the flowering stage. They were then transformed *via* the floral dip method (Clough and Bent, 1998). Seeds harvested from these plants were selected under a fluorescence binocular for expression of the visual marker included in the vectors.

### CRISPR/Cas9 mutagenesis of lncRNAs

To generate deletion mutants, four different sgRNAs for each lncRNA gene were designed using the “CHOPCHOP” website tool (Labun et al., 2016; Labun et al., 2019) to target regions located upstream of the transcription start site in combination with regions within the terminator, to allow complete deletion of the respective gene. sgRNAs were amplified *in vitro*, assembled as previously described (Xie et al., 2015) and cloned into CloneJET (K1231, Thermo Fisher Scientific, Waltham, Massachusetts, US). Each resulting cassette containing the *Arabidopsis* U6-26 promoter, the tRNA complex with four sgRNAs and the pol III terminator was cloned via the *MluI* restriction site into the pDEECO vector (Bente et al., 2020), which contains the egg cell-specific promoter EC1.2p, the *Arabidopsis* codon-optimized Cas9 ORF and the seed-specific GFP marker (Shimada et al., 2010). Transgenic seeds were selected by their green fluorescence and grown into T2 plants. These were genotyped by PCR for the intended deletion and those with homozygous mutant alleles grown into T3 populations. Sequences of sgRNAs and primers used for genotyping are listed in Supplemental Table 2.

### DNA and RNA extraction

DNA for genotyping was obtained by grinding young leaves with glass beads in 400 μl extraction buffer (200 mM Tris pH 8, 250 mM NaCl, 25 mM EDTA) for 3 min at 30 Hz in an MM400 homogenizer (Retsch, Düsseldorf, Germany). After centrifugation of the samples, supernatants were transferred into new tubes and DNA precipitated for >1 h on ice with 1 volume of cold isopropanol and 1/10 volume of sodium acetate (3 M, pH 5.2). Samples were centrifuged 10 min at 16 000 g, pellets washed once in 75% EtOH, air-dried and dissolved in 75 μl H_2_0. PCR reactions were performed with 1.5 μl DNA per sample.

Total RNA was extracted from 14 d-old seedlings using TRI Reagent (Zymo Research) according to the supplier’s protocol. RNA integrity was controlled by electrophoresis on 1.8% agarose-TAE gels. Samples were then treated with Turbo DNase (Invitrogen) according to manufacturer’s instructions. First-strand cDNA synthesis was performed on DNA-free RNA with random hexamer primers and/or (for NATs) gene-specific primers using RevertAid H Minus Reverse Transcriptase (EP0451, Thermo Fisher Scientific, Waltham, Massachusetts, US) or Superscript IV (Invitrogen) according to manufacturer’s recommendations. Absence of gDNA contamination was controlled after 40 PCR cycles on DNA-free RNA and cDNA with primers spanning the intron of the reference gene *AtSAND* (*At2g28390*). Quantitative RT-PCR was performed on a LightCycler96 system (Roche) with FastStart Essential DNA Green Master kit (Roche; Rotkreuz, Switzerland) with ~3 ng of cDNA and three technical replicates. A two-step protocol was run with pre-incubation at 95°C for 10 min followed by 45 cycles at 95°C for 10 sec, 60°C for 30 sec. A final melting cycle at 97°C was done preceding the melting curve analysis. Primer efficiencies were evaluated on a standard curve using a 2-fold or 10-fold dilution series of the samples over 4 dilution points. Relative expression was calculated according to the ΔΔCt method (Livak and Schmittgen, 2001) and normalized to the internal reference genes *ACTIN2* (AT3g18780) or *SAND* (AT2g28390). Relative expression was calculated relative to the WT mock control. Statistical analyses were performed applying Welch Student’s test (α =0.05). Primers are listed in Supplemental Table 2.

### RAPID amplification of cDNA ends (RACE)

Rapid amplification of cDNA ends was performed using the SMARTer^R^ RACE 5’/3’kit (Takara), us 1 μg of DNA-free RNA from zeocin-treated samples as template for first-strand cDNA synthesis according to manufacturer’s instructions. cDNAs were then diluted 2.5 times with Tricine-EDTA buffer and PCR-amplified with gene-specific primers and universal primers UPM (provided in the kit) with the following program: 98°C for 2 min; 35 cycles of 94°C for 30 sec, 65°C for 30 sec and 72°C for 3 min; followed by a final elongation step at 72°C and cooling to 4°C. PCR reactions were run on 1.5% agarose-TAE gels, purified using NucleoSpin Gel and PCR Clean-Up Kit (Takara), cloned into pRACE vector (provided in the kit) and transformed into Stella cells. Between 10-15 colonies were PCR-screened for inserts and the DNA analyzed by Sanger sequencing. Primers are listed in Supplemental Table 2.

### Northern blot

Ten to 20 μg of total RNA were separated on 1.5% GB agarose gel (10 mM Na_2_HPO_4_, 8.4 mM NaH_2_PO_4_, pH 7), blotted onto Hybond NX nylon membrane (Amersham ref. RPN203T) and cross-linked in a UV Stratalinker 2400 (Stratagene, La Jolla, California, US) in auto-crosslink mode. Probes were generated by PCR amplification of the DNA region of interest and labeled through Klenow reaction with [α^32^P]dCTP, using the Amersham Rediprime II Random Prime Labeling System (RPN 1633; GE Healthcare, Chalfont St Giles, UK). Membranes were hybridized (250 mM Na_2_HPO_4_, 7% SDS, 1 mM EDTA, pH 7) overnight at 42°C, followed by washing twice in 2X SSC, 2% SDS solution for 10 min at 50°C. After that, membranes were exposed to a phosphoscreen for 24 h that was scanned on a phosphoimager (Typhoon FLA 9500, GE Healthcare, Chalfont ST Giles, UK). Primers and oligonucleotides used for probe synthesis are listed in Supplemental Table 2.

### RNA sequencing experiments

For sequencing, total RNA was extracted and prepared as described above, with three (UV) or five (zeocin) independent biological replicates per genotype and per condition. Ribosomal RNAs (rRNAs) were removed using Ribo-Zero rRNA removal kit (Illumina). rRNA-free RNAs were controlled on a Fragment Analyzer (Agilent formerly Advanced Analytical, Santa Clara, California, US) with the HS NGS Fragment Kit (DNF-472-0500 RNA, Agilent formerly Advanced Analytical, Santa Clara, California, US). Libraries were prepared with the NEBNext Ultra Directional RNA library Prep kit for Illumina (New England Biolabs, Ma, USA) and sequenced by high-throughput sequencing of pair-end 50 (PE50) by the Next Generation Sequencing Facility (Vienna BioCentre Core Facilities). Details about the sequencing are listed in Supplementary Table 1.

### lncRNA and mRNA data analyses

Raw reads from RNA sequencing were first cleaned (Phred quality score ≥ 20) and trimmed using Trim Galore (Version 0.6.2) in paired-end mode. Read quality was then controlled using FastQC (Version 0.11.8). Processed reads were mapped to the TAIR10 *A. thaliana* reference genome (Lamesch et al., 2012) using STAR (Version 2.7.1a) with the following options: twopassMode Basic, outFilterMultimapNmax 10, alignIntronMax 10000, alignMatesGapMax 6000. Mapped reads from both treated and non-treated samples were then merged and assembled into a unified transcriptome file using Stringtie (Version 2.1.5) (Pertea et al., 2015) with the following options: rf, m = 200, c = 1, s = 2, j = 2.5, f = 0.5 and a = 15 with strand-specific awareness. The assembled transcriptome file was then annotated using gffcompare program with the Araport11 annotation (Cheng et al., 2017) (Version 0.12.1). Protein-coding transcripts and annotated lncRNAs were identified. The remaining unknown transcripts were subjected to further analyses according to the following criteria. 1) Transcripts classified with code “u” (intergenic transcripts), “x” (exonic overlap with reference on the opposite strand), and “i” (transcripts entirely within a reference intron) were retained. 2) Transcripts with low abundance (FPKM max (maximum expression of a lncRNA from all samples) <1) were removed. 3) Transcripts with protein-coding potential were ignored. Protein-coding potential was evaluated using CPC2 (Coding Potential Calculator, CPC >0) (Kang et al., 2017), coding-non-coding index (CNCI, score >0) (Guo et al., 2019), and blastX search against all protein sequences in the Swiss prot database and unannotated with an E-value cut-off >10^-4^. Reads overlapping “transcripts” features in the assembled transcriptome file were counted using the FeatureCounts function from Subread package (Version 2.0.1). Differential gene expression analysis was estimated with DESeq2 (https://bioconductor.org). In any pairwise comparison, lncRNAs or mRNAs with a filter of adjusted p-value <0.05 and absolute fold change of 1.5 were considered as differentially expressed. R and Bioconductor (https://bioconductor.org) were used to plot data. Details about transcript assembly and differentially expressed genes are listed in Supplementary Tables 3 and 4.

### Conservation analysis within Arabidopsis accessions

SNP numbers were determined with the SNP-calling data from the 1001 Genomes Genome Consortium (https://1001genomes.org/data/GMI-MPI/releases/v3.1/1001genomes_snp-short-indel_only_ACGTN.vcf.gz) using vcftools (v.0.1.16) to extract SNP positions in the TAIR10 genome and mapBed (bedtools v.2.27.1) to count the SNP number for each locus, followed by normalization by the locus length. Fully assembled genomes of 27 accessions (Col-0 and 26 non-reference accessions) were provided by the Nordborg lab, GMI, Austria. For each lncRNA, the sequence corresponding to the transcript and the surrounding 300 bp up- and downstream was extracted from the TAIR10 genome and blasted onto the 27 genomes using blastn (blast+ v2.8.1) with the following options: -word_size 10 -strand both -outfmt 7 -evalue 1e-7. The multiple sequence alignment was obtained and displayed using Unipro UGENE v43.0 “Align with Muscle” option. To find the gene copies, the blastn results were filtered for sequences with >80% sequence identity and >80% length match to the TAIR sequence, allowing for insertions of up to 1.5 kb to account for possible TE insertions in the non-reference accession genomes. For gene expression calculation, we used RNA-seq data from mature leaves (Kawakatsu et al., 2016, GEO accession number GSE80744) and RNA-seq data from 7-day-old seedlings, 9-leaf rosettes, flowers (with flower buds), and pollen (Kornienko et al., 2023, GEO accession number GSE226691). Raw RNA-seq reads were mapped to the TAIR10 genome using STAR (v.2.7.1) and exonic read counts were calculated using feature counts software from the subread package (v.2.0.0). Raw reads were normalized by transforming them into TPMs. Expression variability was calculated as coefficient of variance: standard deviation of expression across accessions divided by the mean expression level. Admixture groups (geographic origin) information for different accessions was obtained from the 1001 Genomes Genome Consortium (https://1001genomes.org/data/GMI-MPI/releases/v3.1/1001genomes_snp-short-indel_only_ACGTN.vcf.gz).

### Phylogenetic analysis within the Brassicaceae

For an initial overview over the presence of the lncRNA B, C and D, genomes in the EnsEMBL plant data base were screened for homologs using BLASTN. Since there were only hits within the Brassicaceae, homologous sequences of specific Brassicaceae genomes were obtained from EnsEMBL plant and two additional genome databases, CoGe and NCBI Genomes (via NCBI Taxonomy), using the respective online BLAST interfaces. In all interfaces, the most sensitive BLAST mode offered was applied, namely “Distant homologies against the Genomic sequence” in EnsEMBL Plants, “Somewhat similar sequences (blastn)” in NCBI Genomes and “E-value cut-off: 1, blastn matrix 1-2, Gap Penalties: 5 2” on the CoGe (Comparative Genomics) website (https://genomevolution.org/coge/). Because the lncRNAs have regions prone to being filtered out by low-complexity filters, reducing the probability to find hits, low-complexity filtering was switched off where possible for both query and genome.

Hits were called significant if they had E-values of at least 10^-4^ and had a length of at least 100 nt, or if they had two non-overlapping hits within the query with E-values of at least 10^-4^ summing up to a total length of at least 100 nt. Sequences were extracted from the genomes from 300 nt downstream to 300 nt upstream of the total hits using samtools faidx from the SamTools package (Danecek et al., 2021, v. 1.15).

The sequences were aligned with MAFFT (Katoh and Standley, 2016, v. 7.487) using accurate options “--reorder --maxiterate 1000 --localpair”. In a quality control step, some sequences not showing sufficient similarity along the query to produce an unambiguous alignment had to be discarded as false-positives possible found due to random matches, e.g., in low-complexity regions. Subsequently, the remaining sequences were aligned again with the above parameters and then pruned to the full-length transcripts obtained in the Arabidopsis experiments.

The final alignments served as input for a phylogenetic tree reconstruction using IQ-TREE (Minh et al., 2020, v. 2.1.3) with 10,000 UFboot (Minh et al., 2013) samples using the parameters “-keep-ident-bb 10000”. The best-fit models of evolution were obtained by ModelFinder as implemented in IQ-TREE (Kalyaanamoorthy et al., 2017) using BIC.

## Acknowledgements

The authors are very grateful to Arndt von Haeseler (Center for Integrative Bioinformatics Vienna (CIBIV)) for constructive comments on the manuscript. They want to thank the Plant Science and Next Generation Sequencing Departments of the Vienna BioCenter Core Facilities (VBCF), as well as the Molecular Biology Services, the Media Preparation Lab, the Environment/Health/Safety Group, and the Lab Support of the GMI/IMBA/IMP, for their excellent support. N.D. is grateful for a Lise Meitner Fellowship from the Austrian Science Fund (M2410).

## Figure legends

**Figure S1:**
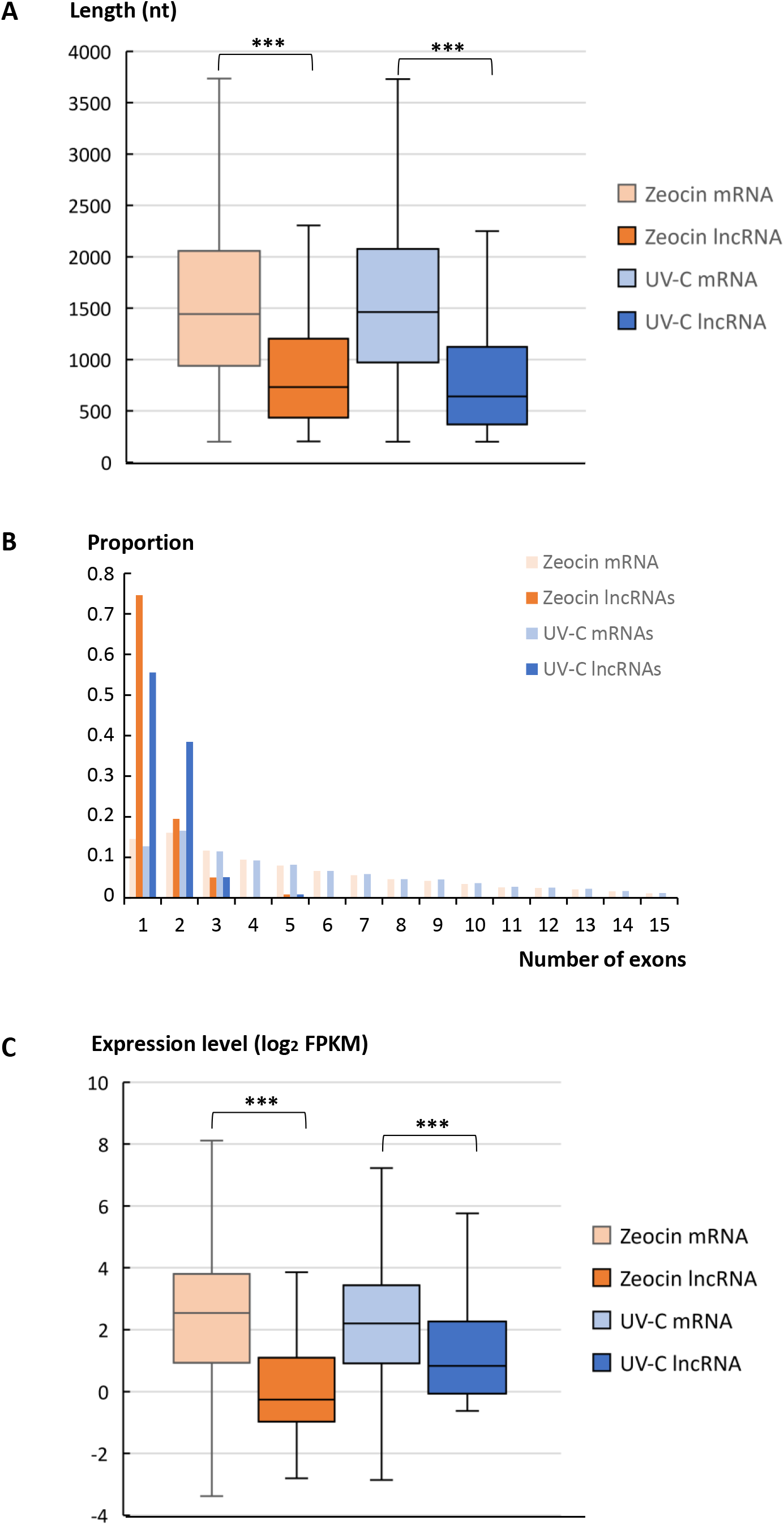
Characterization of novel lncRNAs. (A) Exon number distribution in genes for mRNAs and lncRNAs present in libraries from zeocin-or UV-C-treated samples. (B) Length distribution of mRNAs and lncRNAs present in libraries from zeocin-or UV-C-treated samples. (p-value <0.000001, Mann-Whitney test). (C) Expression levels (log2 FPKM) of mRNAs and lncRNAs present in libraries from zeocin-or UV-C-treated samples. (p-value <0.000001, Mann-Whitney test).

**Figure S2:**
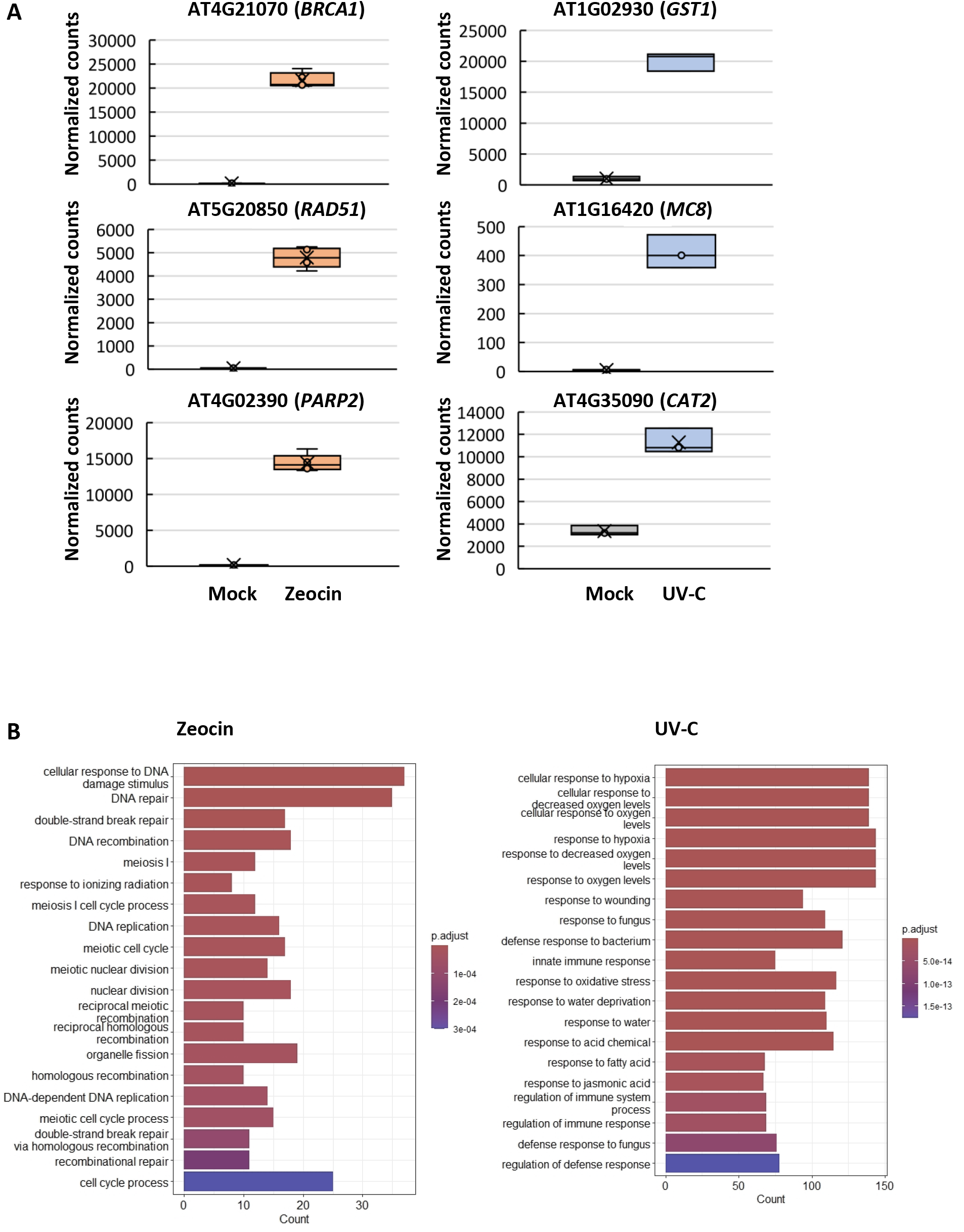
Validation of DNA damage induction. (A) Normalized counts for genes with known functions in DNA repair: *BRCA1* (AT4G21070), *RAD51* (AT5G20850), *PARP2* (AT4G02390) in zeocin-treated samples; *GST1* (AT1G02930), *MC8* (AT1G16420), *CAT2* (AT4G35090) in UV-C-treated samples. Error bars indicate standard deviation of 3 biological replicates (*Welcher test* **p-value <0.01, *p-value <0.05). (B) GO-term enrichment of differentially expressed genes in zeocin-or UV-C-treated samples.

**Figure S3:**
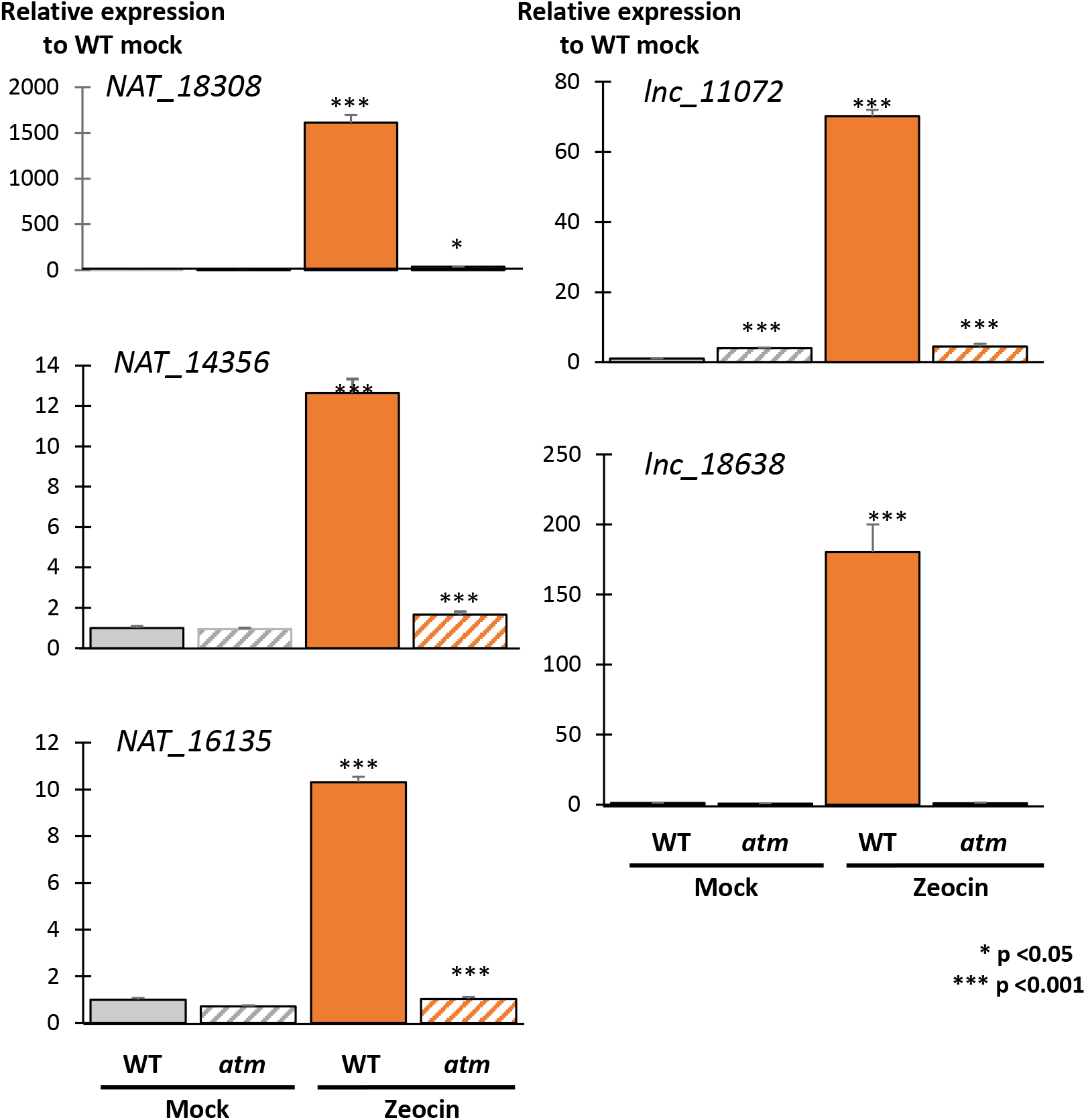
Induction of other lncRNAs by zeocin. Expression of additional novel lncRNAs in WT or *atm* mutant in mock-or zeocin-treated samples, normalized to a constitutively expressed actin gene. Data were normalized to the values in WT mock samples. Error bars indicate standard deviation of 3 biological replicates (*Welcher test* **p-value <0.01, *p-value <0.05).

**Figure S4:**
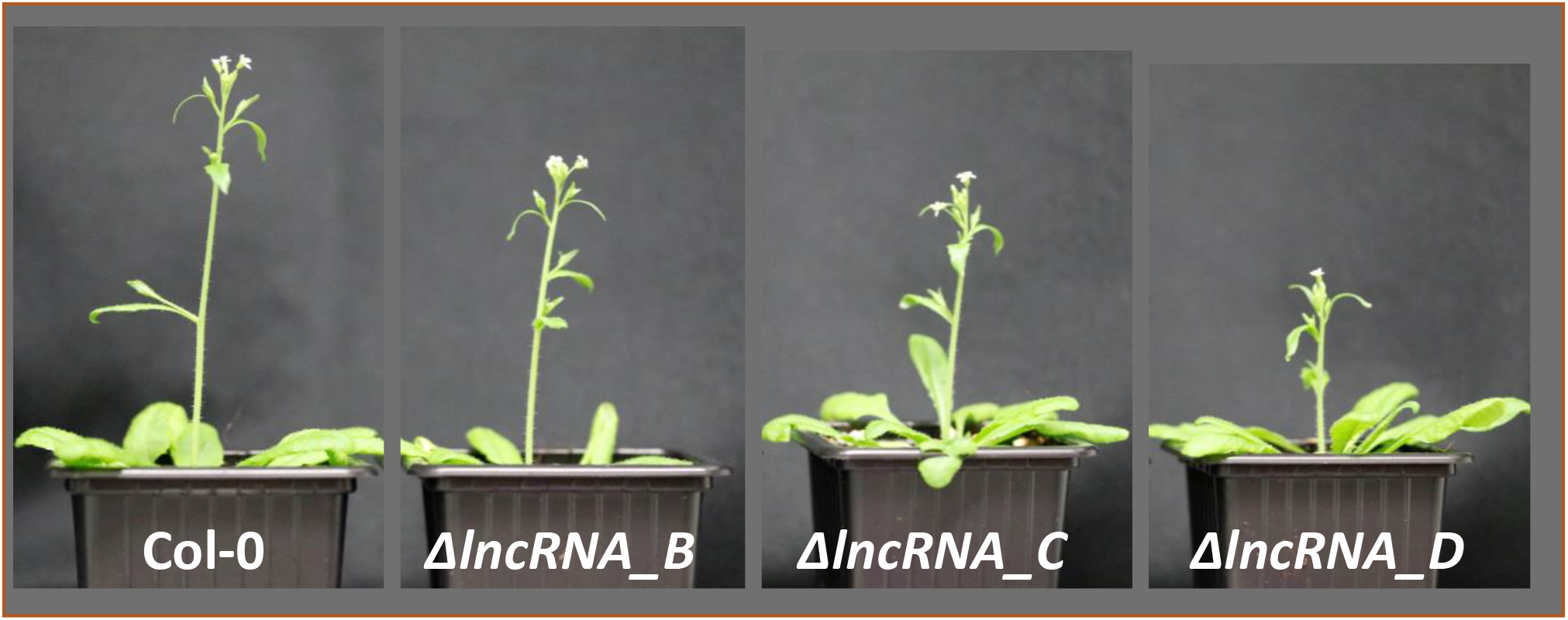
Phenotype of deletion mutants. Comparison between WT (Col-0, left) and mutants lacking lncRNA B, C, or D, for soil-grown plants at the flowering stage.

**Figure S5:**
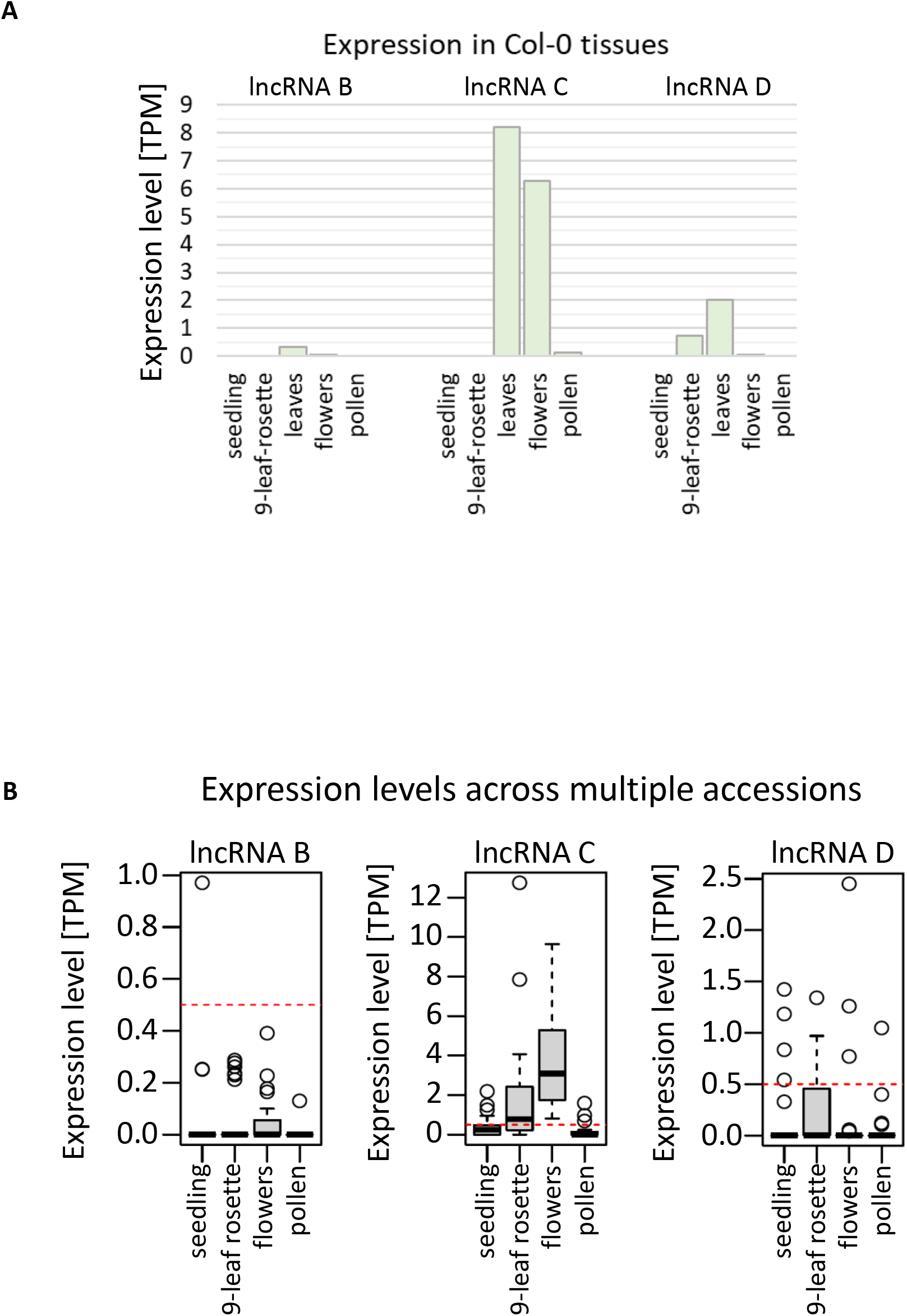
lncRNA expression in tissues and across accessions. (A) Expression in different tissues of the reference accession Col-0. (B) Expression in different tissues across multiple accessions. The red dashed horizontal line indicates expression cut-off (TPM=0.5). Expression is calculated from RNAseq data as in Figure 5 D.

***Table S1: Summary of sequencing data***

Tables show the number of processed reads, the percentage of reads uniquely mapped to the genome, and the percentage of reads assigned to the assembly for zeocin- and UV-C-treated samples.

***Table S2: Sequences of primers and gRNAs***

The table lists the names, sequences, and applications of oligonucleotides used in the study.

***Table S3: Transcript assembly***

The tables list mRNAs, known lncRNAs, and novel lncRNAs assembled in zeocin- and UV-C-treated samples, including their chromosomal location, genome coordinates, and read frequencies.

***Table S4: List of differentially expressed genes***

The tables list genes that are differentially expressed between mock control and exposure to either zeocin or UV, including their chromosomal location and their gene product type.

